# *TractoInferno*: A large-scale, open-source, multi-site database for machine learning dMRI tractography

**DOI:** 10.1101/2021.11.29.470422

**Authors:** Philippe Poulin, Guillaume Theaud, Francois Rheault, Etienne St-Onge, Arnaud Bore, Emmanuelle Renauld, Louis de Beaumont, Samuel Guay, Pierre-Marc Jodoin, Maxime Descoteaux

**Affiliations:** Sherbrooke Connectivity Imaging Laboratory (SCIL), Université de Sherbrooke, Quebec, Canada; Videos and Images Theory and Analytics Laboratory (VITAL), Université de Sherbrooke, Sherbrooke, Canada; Montreal Sacred-Heart Hospital Research Centre, Montréal, QC, Canada; Department of Surgery, Université de Montréal, Montréal, QC, Canada

**Keywords:** Neuroimaging, Database, Tractography, Machine Learning

## Abstract

*TractoInferno* is the world’s largest open-source multi-site tractography database, including both research- and clinical-like human acquisitions, aimed specifically at machine learning tractography approaches and related ML algorithms. It provides 284 datasets acquired from 3T scanners across 6 different sites. Available data includes T1-weighted images, single-shell diffusion MRI (dMRI) acquisitions, spherical harmonics fitted to the dMRI signal, fiber ODFs, and reference streamlines for 30 delineated bundles generated using 4 tractography algorithms, as well as masks needed to run tractography algorithms. Manual quality control was additionally performed at multiple steps of the pipeline. We showcase *TractoInferno* by benchmarking the learn2track algorithm and 5 variations of the same recurrent neural network architecture. Creating the *TractoInferno* database required approximately 20,000 CPU-hours of processing power, 200 man-hours of manual QC, 3,000 GPU-hours of training baseline models, and 4 Tb of storage, to produce a final database of 350 Gb. By providing a standardized training dataset and evaluation protocol, *TractoInferno* is an excellent tool to address common issues in machine learning tractography.

## 1. Introduction

Tractography is the computerized process of reconstructing brain white matter fibers from diffusion MRI (dMRI) data. It usually consists of three steps : i) estimating local fiber directions from carefully pre-processed diffusion-weighted images (DWI) (e.g. denoising, eddy, motion, susceptibility corrections), ii) reconstructing white matter pathways (i.e. tractography), and iii) delineating bundles (a group of similar streamlines connecting the same brain regions) [1, 2].

Current “traditional” tractography approaches (deterministic and probabilistic) mostly rely on making local point-wise decisions in the fiber ODF field, iterating until termination [3, 4]. Global methods have also been proposed [5, 6, 7, 8], but Rheault et al. mentions that “[…] global tractography methods ultimately rely on local information patched together” and “even global tractography algorithms struggle to correctly assemble a streamline” [9]. Tractogram filtering [10, 11, 12, 13] is also a popular post-processing method used to remove streamlines that do not fit anatomical constraints (such as explaining the underlying signal), but requires an over-complete tractogram as it does not create new streamlines, thus effectively “wasting” computing power. Finally, streamline clustering [14, 15] can be used to group streamlines based on similarity and remove possible outliers, but it suffers from the same drawback as tractogram filtering, as it requires an over-complete tractogram.

These approaches mostly rely on mathematical models or anatomical priors, and do not require histological ground truth to work. However, this is an issue for machine learning algorithms, where the training dataset is an integral part of the resulting model [16]. Machine learning methods need reference streamlines to train on. Unfortunately, on real datasets, streamlines can only be generated by traditional [and yet non-machine learning] tractography methods, which are imperfect by their very nature [2]. This is an issue for testing if the predictions made by these methods are reliable or not. Luckily, by combining streamlines (both true positives and false positives) generated by several tractography algorithms and using filtering and clustering to remove as much false positives as possible, it is possible to establish a *gold standard* reference dataset. Even without a histologically accurate ground truth, it would be desirable to have algorithms that can reproduce a gold standard reference while generating as little false positive streamlines as possible.

In the recent years, machine learning algorithms have been proposed to improve the second step of the process by some combination of 1) taking advantage of the full diffusion information or other modalities, 2) generating more reliable streamlines using a reference teacher dataset, or 3) integrating more spatial context to guide the tracking process (either neighbourhood or path information) [16, 17, 18, 19, 20]. For example, *TractSeg* [19] is a method that first identifies the volume of reference of a specific white matter bundle, and then generates a bundle-specific tractogram by running a traditional tractography algorithm inside the bundle mask only. To do so, convolutional neural networks [21] learn to map the diffusion volume to multiple binary bundle segmentation maps. *LearnToTrack* [18] and *DeepTract* [22] propose to use information along a streamline path to guide its generation process (instead of making point-wise decisions) using Recurrent Neural Networks [23, 24]. *Entrack* [20] proposes a Neural Network with a fixed context of 4 streamline steps, and models a probabilistic streamline direction using a von Mises-Fisher distribution trained with entropy regularization.

Unfortunately, these machine learning methods train and evaluate their models on different dataset which makes it difficult to compare their true generalization capabilities [16]. It is often a combination of the ISMRM2015 Tractography Challenge [2] and some subjects from the HCP Young Adults database [25]. Additionally, data pre-processing might vary between proposed methods, and different algorithms and protocols are used to generate the reference tracts. Finally, evaluating the generalizability of a model is almost impossible without diverse (aka multi-site) training and test sets. As a result, all those discrepancies in methodology make it very challenging to assess the reliability of a single approach, and make it almost impossible to fairly compare approaches.

We propose to address this problem by building *TractoInferno*: the largest publicly available, multi-site, dMRI and tractography database, which provides a new baseline for training and evaluating machine learning tractography methods. It provides 284 datasets acquired from 3T scanners across 6 different sites. *TractoInferno* includes T1-weighted images, single-shell diffusion MRI (dMRI) acquisitions, spherical harmonics fitted to the dMRI signal, fiber ODFs, and reference streamlines for 30 delineated bundles generated by combining 4 different tractography algorithms, as well as masks needed to run tractography algorithms.

We use *TractoInferno* to benchmark the 4 tractography algorithms used to create the reference tractograms, along with the learn2track [18] algorithm and 5 variations of the same recurrent neural network architecture, inspired in part by the models of (Benou and Riklin Raviv) and (Wegmayr and Buhmann) [22, 20].

Creating the *TractoInferno* database required approximately 20,000 CPU-hours of processing power, 200 man-hours of manual QC, 3,000 GPU-hours of training baseline models, and 4 Tb of storage, to produce a final database of 350 Gb.

*TractoInferno* is a dataset intended to promote the development of machine learning tractography algorithms, which generally suffer from multiple issues, such as limited datasets or inconsistent training data. Its large-scale and multi-site aspect is an undeniable benefit to best evaluate the generalization capabilities of new ML algorithms. We consider *TractoInferno* to be by far the best available tool for training, evaluating, and comparing future machine learning tractography algorithms.

## 2. Datasets

The proposed dataset is made of a combination of six dMRI databases, either publicly available and free to redistribute or acquired through open-access data sharing agreements. Databases were chosen with the explicit goal of having a diversity of scanner manufacturers, models, and protocols. We chose to fix certain parameters for uniformity, such as having only 3T scanners, and b-values of around 1000 s/mm^2^, as we don’t know how they could affect machine learning models. The focus is effectively on assessing the reliability of algorithms under different scanner manufacturers and acquisition protocols. We obtained an initial number of data from 354 subjects, with the original metadata described in Table 1.

**Table 1:** Original datasets metadata. Not all metadata information was available from the original datasets. Missing metadata is reported as {N/A}. Resolution is in mm^3^ isotropic. b-value is in s/mm^2^. TR and TE are in ms. *: 21 directions acquired twice by reversing the gradient polarity, then averaged over another identical acquisition (total of 84 DWI volumes). **: 64 directions acquired twice, not averaged.

| Name | *Mazoyer*<br>*et. al*<br>*[26]* | *Tsushida*<br>*et. al*<br>*[27]* | *DeLuca*<br>*et. al*<br>*[28]* | *Poldrack*<br>*et. al*<br>*[29]* | *Tamm*<br>*et. al*<br>*[30]* | *Tremblay*<br>*et. al*<br>*[31]* |
| --- | --- | --- | --- | --- | --- | --- |
| Scanner | 3T<br>Philips<br>Achieva | 3T<br>Siemens<br>Prisma | 3T<br>Siemens<br>Prisma | 3T<br>Siemens<br>Trio | 3T GE<br>Discovery<br>MR750 | 3T<br>Siemens<br>Magnetom<br>TIM Trio |
| # subjects | 39 | 20 | 64 | 130 | 86 | 15 |
| Age avg | 28.1 | 21.4 | 31.9 | 31.3 | N/A | 58.1 |
| Age std | 7.3 | 1.7 | 7.6 | 8.7 | N/A | 5.3 |
| F/M | 0/39 | 10/10 | 49/15 | 62/68 | 44/42 | 0/15 |
| L/R | 8/31 | N/A | 0/64 | N/A | N/A | 3/12 |
| Resolution | 2 | 1.75 | 2 | 2 | 2.3 | 2 |
| b-value | 1000 | 1000 | 1000 | 1000 | 1000 | 700 |
| TR | 8500 | 3540 | 1800 | 9000 | 7000 | 9200 |
| TE | 81 | 75 | 70 | 93 | 81 | 84 |
| Nb dirs | 21* | 32 | 128** | 64 | 45 | 30 |

### 2.1. Mazoyer et. al - BIL & GIN

We retained 39 subjects from the BIL&GIN database [26], acquired on a 3T Philips Achieva, with the following dMRI protocol: TR = 8500 ms, TE = 81 ms, angle = 90°, SENSE reduction factor = 2.5, FOV 224 mm, acquisition matrix 112 × 112, 2 mm^3^ isotropic voxel.

The dMRI acquisition consisted of 21 gradient directions at b = 1000 s/mm^2^, acquired twice by reversing the polarity, and then repeated twice for a total of 84 DWI images, averaged down to a single volume with 21 directions. A single b = 0 s/mm^2^ was also acquired alongside the DWI images. Subjects were all males, with age mean/std of 28.1 +− 7.3 (Min: 20, Max: 57). 8 subjects were left-handed and 31 right-handed.

### 2.2. Tsushida et. al - MRi-Share

We obtained 20 subjects from the MRi-Share database [27], acquired on a 3T Siemens Prisma, with a dMRI protocol designed to emulate the UKBioBank project [32], specifically: TR = 3540 ms, TE = 75 ms, 1.75 mm^3^ isotropic voxel.

We selected the b = 1000 s/mm^2^ DWI images only, consisting of 32 gradient directions, and 3 provided b = 0 s/mm^2^ images. Subjects were composed of 10 females, 10 males, with age mean/std of 21.4 +− 1.7. Minimum/maximum age and handed-ness metadata were not available.

### 2.3. DeLuca et. al - Bilingualism and the brain

We have 64 subjects from the *Bilingualism and the Brain* database [28, 33], acquired on a 3T Siemens Prisma, with the following dMRI protocol: Echo planar imaging, TR = 1800 ms, TE = 70 ms, acquisition matrix 256 × 256, 2 mm^3^ isotropic voxel.

The dMRI acquisition consisted of 64 gradient directions at b = 1000 s/mm^2^, acquired twice, and 4 b = 0 s/mm^2^ images. Subjects were composed of 49 females and 15 males, with age mean/std of 31.9 +− 7.6 (Min: 18, Max: 52). All subjects were right-handed.

### 2.4. Poldrack et. al - UCLA CNP

We got 130 healthy subjects from the *UCLA Consortium for Neuropsy-chiatric Phenomics LA5c Study* [29], acquired on a 3T Siemens Trio, with the following dMRI protocol: echo planar imaging, TR = 9000 ms, TE = 93 ms, acquisition matrix 93 × 93, 90 degree flip angle, 2 mm^3^ isotropic voxel. DWI were corrected for eddy currents and head motion using the b0 images as reference.

The dMRI acquisition consisted of 64 gradient directions at b = 1000 s/mm^2^, and 1 b = 0 s/mm^2^ image. Subjects consisted of 62 females and 68 males, with age mean/std of 31.3 +− 8.7 (Min: 21, Max: 50). Handed-ness metadata was not available.

### 2.5. Tamm et. al - The Stockholm Sleepy Brain Study

We retained 86 subjects from the Stockholm Sleepy Brain Study database [30, 34], acquired on a 3T GE Discovery MR750, with the following dMRI protocol: Echo planar imaging, TR = 7000 ms, TE = 81 ms, 2.3 mm^3^ isotropic voxel.

The dMRI acquisition consisted of 45 gradient directions at b = 1000 s/mm^2^, along with 5 b = 0 s/mm^2^ images. Subjects were composed of 44 females and 42 males, with 47 subjects in the [20-30] years old bracket and 39 subjects in the [65-75] years old bracket. Handedness was not available.

### 2.6. Tremblay et. al - mTBI and Aging study (controls)

We obtained 15 subjects from the mTBI and Aging Study [31], all controls from the “remote” group. they were acquired on a 3T Siemens Magnetom TIM Trio, with the following dMRI protocol: TR = 9200 ms, TE = 84 ms, 2 mm^3^ isotropic voxel.

The dMRI acquisition consisted of 30 gradient directions at b = 700 s/mm^2^. along with 1 b = 0 s/mm^2^ image. Subjects were all males, with age mean/std of 58.1 +− 5.3 (Min: 52, Max: 67). 3 subjects were left-handed and 12 were right-handed.

## 3. Methodology

We processed the original acquisition volumes of the 354 aforementioned subjects with the same pipeline to offer a uniform database of dMRI images, derivatives, and bundle tractograms. First, all original DWI went through a manual quality control (QC) step to remove any obvious errors prior to the processing pipeline. Then, the *TractoFlow* pipeline was run to process the data and compute necessary derivatives [35, 36, 37]. Another QC step was executed afterwards, to remove images with artifacts that could not be corrected automatically. Next, ensemble tractography was performed using four different algorithms to extract a diverse set of streamlines: deterministic tractography [38], probabilistic tractography [39], Particle-Filtered Tractography [40] and Surface-Enhanced Tractography [41]. RecoBundlesX (RBX) was used subsequently to perform bundle extraction on the whole-brain tractograms, using the default suggested bundle models [42, 43]. A final manual QC step was performed to examine the extracted bundles, and remove anything that contained obvious mistakes, or did not meet our criteria for bundle extraction. All manual quality control steps were done using dmriqcpy (https://github.com/scilus/dmriqc_flow). Figure 1 shows the processing steps of *TractoInferno*.

**Figure 1:**
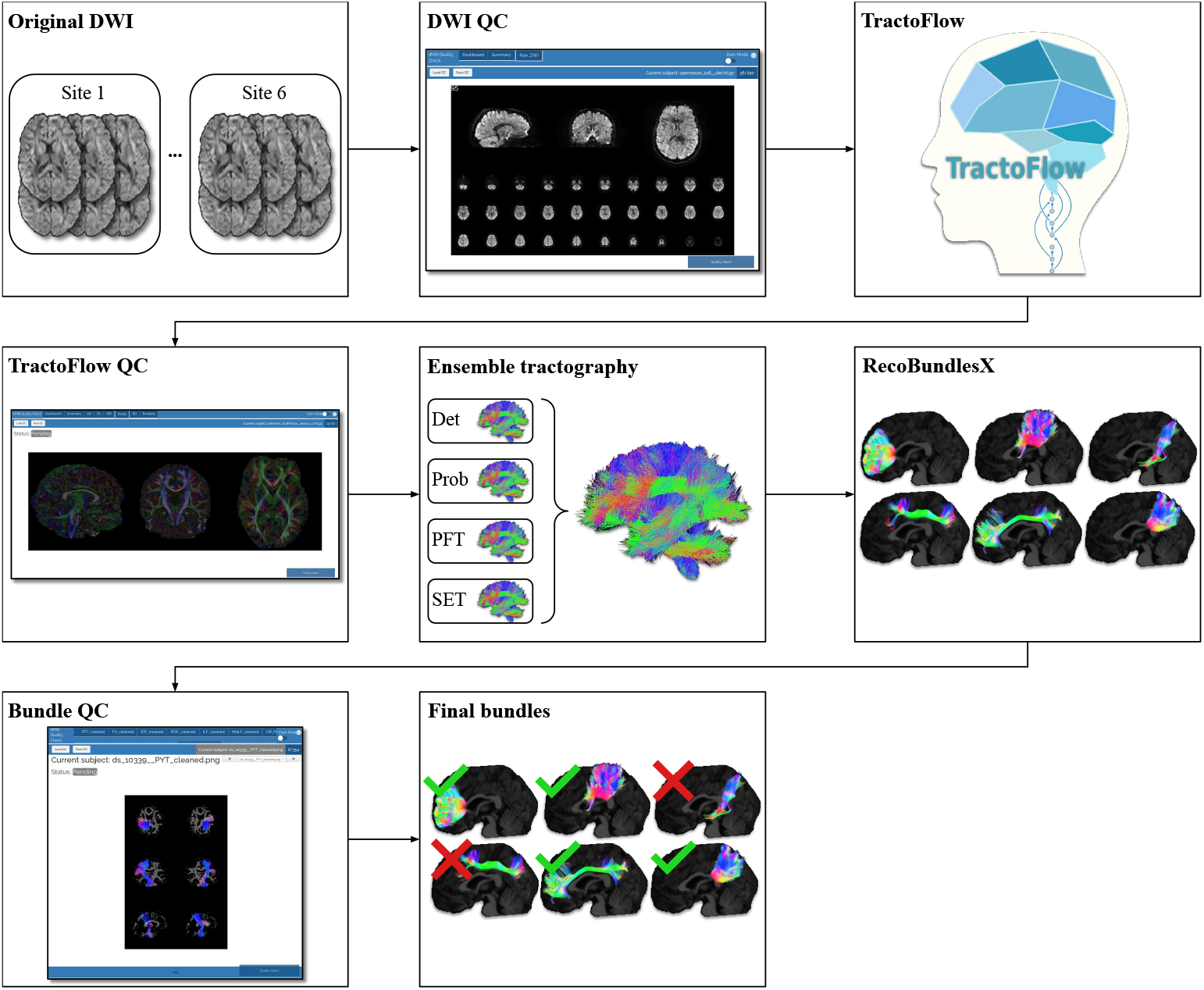
*TractoInferno* processing pipeline, from original DWI images to final bundles.

From the initial 354 volumes, after all the processing steps and quality control, we were left with 284 volumes and associated bundles. The final volumes were split into training, validation and test sets with a 70%/20%/10% split for reproducibility across future experiments. The specific commands for the whole pipeline are available in Appendix A. For a final dataset size of 350Gb, we needed approximately 20,000 CPU-hours of processing time, 200 man-hours of manual QC, and 4 Tb of storage. The models bench-marked in section 5 also required an additional 3,000 GPU-hours for training and generating candidate tractograms. In the next sub-sections, we detail the TractoInferno processing steps.

### 3.1. Raw data QC

We used *dmriqcpy* to generate QC reports. These reports are in HTML format so it is easily assessed and annotated by multiple people. The raw data reports contain multiple tabs with complementary information, as shown in Figure 2. Three different raters went through the QC reports and individually rated every acquisition with a “score” (either *pass, fail*, or *warning*) and comment if necessary. Specifically, failure cases included the presence of visual artifacts (e.g. missing slices, low signal-to-noise ratio, corrupted data, high spatial distortion) and other artifacts harder to identify (such as a “broken” gradient acquisition scheme). Afterwards, all subjects tagged as “fail” were removed, and considered as impossible to repair with our available tools. All subjects tagged as “pass” or “warning” were passed on for TractoFlow, the next step in the pipeline. Subjects tagged as “warning” were re-examined after the TractoFlow processing to examine if any issues remained, or if they were compensated for by the pipeline.

**Figure 2:**
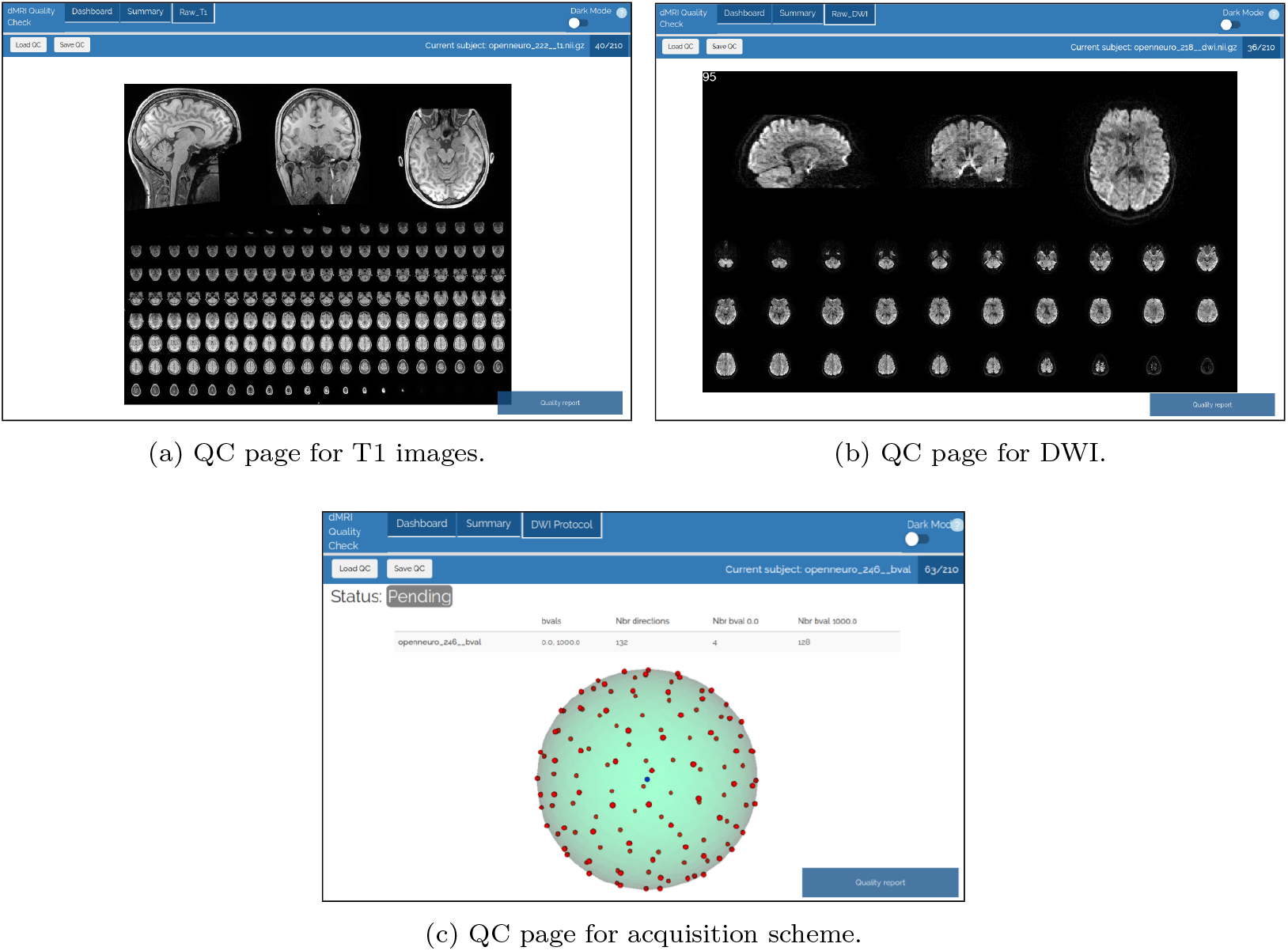
Examples of HTML pages generated by dmriqcpy for data QC. (a) 3 slices of the T1 image (one for each axis), plus a mosaic of multiple axial slices. (b) 3 GIFs of the dMRI (one slice in each axis), plus a mosaic of multiple axial slices; (c) The gradient directions represented on a sphere.

### 3.2. TractoFlow pipeline

We used TractoFlow 2.1.1 [35] to process the raw DWI. To make sure that every processing step was traceable and reproducible, a Singularity [36] image was used along with the Nextflow pipeline [37]. Note however that some results may not be 100% reproducible due to the uncertain nature of registration, parallel processing, and floating point precision. We ran the full pipeline except for the *Topup* process, as not all reverse b0 images were available [44]. Specifically, the pipeline executed the following steps:

- DWI brain extraction [45], denoising [46], eddy current correction [47], N4 bias field correction [48], cropping, normalization [49, 50], and resampling [51];
- T1 denoising [52], N4 bias field correction [48], registration [53] and tissue segmentation [54] maps for Particle-Filtered Tractography [40, 55];
- DTI fitting and metrics extraction [56];
- fODF fitting using constrained spherical deconvolution [57, 58, 59], with a fiber response function fixed manually to [0.0015, 0.0004, 0.0004].

### 3.3. TractoFlow results QC

Outputs from TractoFlow went through a manual QC pass to identify failure cases. Using *dmriqcpy*, we were able to easily and quickly look at all maps derived from DTI and fODF metrics, along with T1 registration overlay. For example, RGB maps extracted from DTI metrics allowed us to quickly identify if tensor peaks were well-aligned or if a flip was needed, and T1 registration overlays showed whether too much deformation was present.

### 3.4. Ensemble tractography

Using a single tractography method as reference for a machine learning algorithm might induce unwanted biases. To avoid this, we chose to use ensemble tractography by combining 4 different algorithms to generate reference streamlines, namely deterministic [38], probabilistic [39], particle-filtered [40], and surface-enhanced [41] tractography. We fixed the tracking parameters to the standard default values:

- WM + WM/GM interface seeding
- 10 seeds per voxel (Det, Prob, PFT) or 10,000,000 surface seeds (SET)
- Step size 0.2mm (Det, Prob, SET) or 0.5mm (PFT)
- WM tracking mask (Det, Prob) or WM/GM/CSF probability maps (PFT, SET)

We detail each algorithm in the following three subsections.

#### 3.4.1. Deterministic tracking

Deterministic tracking [38] chooses the fODF peak most aligned with the previous direction as the next streamline step. It seems better suited to connectomics studies [3], mainly on account of the low number of false positives it produces. While it may be inadequate for spatial exploration and bundle reconstruction, deterministic tracking essentially produces smooth streamlines that follow the easiest path through the fODF field. Smooth streamlines are likely more desirable for machine learning algorithms rather than chaotic streamlines that often change directions locally.

#### 3.4.2. Probabilistic tracking and Particle-Filtered Tractography

Probabilistic tracking [39] samples a new streamline direction inside a cone of evaluation aligned with the previous direction, with a probability distribution proportional to the shape of the fiber ODF within the cone.

Particle-Filtered Tractography [40] is an improvement over probabilistic tracking. It takes as input probability maps for streamline continuation/stopping criteria, and allows to “go back” a few steps when a streamline terminates in a region not included in the “termination-allowed” map.

Both algorithms are better suited for spatial exploration, at the cost of producing much more false positives. They are especially effective for bundle reconstruction, in which case there are anatomical priors about both the endpoints that should be connected and the pathway that should be followed by the bundle.

#### 3.4.3. Surface-Enhanced Tracking

Finally, Surface-Enhanced Tracking [41] is a state-of-the-art tractography algorithm that relies on initializing streamlines in an anatomically plausible way at the cortex, then running a PFT tracking algorithm. Indeed, gyri have been shown to be problematic regions for tractography, where low dMRI resolution can lead to a gyral bias in streamline terminations [60].

To this end, we computed the WM-GM boundary surface from the T1w image using the *CIVET* [61] tool and the CBRAIN [62] platform. Then, SET uses a geometric flow method, based on surface orthogonality, to reconstruct the fanning structure of the superficial white matter streamlines. The output of this flow is used to initialize and terminate a PFT tractography algorithm. The result is a tractogram with improved cortex coverage, improved fanning structure in gyri, and reduced gyral bias.

### 3.5. Bundle segmentation with RBX

We used RBX [42, 43] to automatically extract WM bundles. The algorithm works by matching streamlines to an atlas of reference bundles. First, a quick registration step brings the atlas into native space using the atlas FA image. Then, a whole-brain tractogram is compared against the bundles atlas using multiple sets of parameters to extract a fixed set of bundles, listed in Table 2. Finally, a majority voting step extracts the final streamlines for each bundle.

**Table 2:** List of bundles in the default RBX atlas.

|  |  |
| --- | --- |
| AC | Anterior commissure |
| AF | Arcuate fasciculus |
| CC_Fr_1 | Corpus callosum, Frontal lobe (most anterior part) |
| CC_Fr_2 | Corpus callosum, Frontal lobe (most posterior part) |
| CC_Oc | Corpus callosum, Occipital lobe |
| CC_Pa | Corpus callosum, Parietal lobe |
| CC_Pr_Po | Corpus callosum, Pre/Post central gyri |
| CC_Te | Corpus callosum, Temporal lobe |
| CG | Cingulum |
| FAT | Frontal aslant tract |
| FPT | Fronto-pontine tract |
| FX | Fornix |
| ICP | Inferior cerebellar peduncle |
| IFOF | Inferior fronto-occipital fasciculus |
| ILF | Inferior longitudinal fasciculus |
| MCP | Middle cerebellar peduncle |
| MdLF | Middle longitudinal fascicle |
| OR_ML | Optic radiation and Meyer's loop |
| PC | Posterior commissure |
| POPT | parieto-occipito pontine tract |
| PYT | Pyramidal tract |
| SCP | Superior cerebellar peduncle |
| SLF | Superior longitudinal fasciculus |
| UF | Uncinate fasciculus |

The whole pipeline was run using a Singularity container [36] and Nextflow [37] for reproducibility. It is freely available online (https://github.com/scilus/rbx_flow/), along with a suggested bundles atlas (https://zenodo.org/record/4630660#.YJvmwXVKhdU).

### 3.6. Bundle segmentation QC

#### 3.6.1. Automated pre-QC

To facilitate the QC procedure, we ran a pre-QC analysis to automatically rate bundles according to pre-defined criteria before manual inspection. These criteria are detailed in Table 3. Afterwards, all bundles were looked at manually through an easier procedure that consists in confirming an already assigned rating rather than rating from scratch.

**Table 3:** Automatic rating criteria, in order of priority. *x* is the number of streamlines of the bundle of interest; *μ* and *σ* are the average and the standard deviation, respectively, of the number of streamlines for the bundle of interest, across all subjects;

| Rating | Criteria |
| --- | --- |
| Fail | $x < 50$<br>$x == 0$ in either hemisphere (if symmetric bundle). |
| Warning | $x \notin [\mu - 1.5\sigma, \mu + 3.5\sigma]$ |
| Pass | $x \in [\mu - 1.5\sigma, \mu + 3.5\sigma]$ |

#### 3.6.2. Manual quality control using dmriqcpy

A bundle was removed if it looked visually incomplete or if it deviated from the expected pathway. A poor bundle reconstruction might have an algorithmic cause, such as sub-optimal tracking parameters or improper registration in RBX. It might also have an anatomical cause, such as unknown or undisclosed neurological conditions. Furthermore, visually evaluating a bundle reconstruction is very subjective, and a rater’s evaluation can be affected by the time of day, duration of QC, or even the angle of visualization in the QC tool [63]. For all those reasons, and with the goal of establishing a gold standard for machine learning tractography methods, we chose to be somewhat severe in the rating of bundles, in order to minimize the number of false positives, even if that meant missing out some true positive data. After QC, we chose to ignore the following bundles from the atlas due to generalized reconstruction errors : AC, CC Te, Fx, ICP, PC, SCP.

## 4. Evaluation pipeline for candidate tractograms

When evaluating machine learning tractography algorithms, we focus on the volume covered by the recognized bundles (compared to the gold standard bundles). We make no assumptions about the ability to “explore” the brain outside the scope of the *TractoInferno* dataset. Consequently, we ignore anything that is not recognized as a candidate bundle, and do not try to categorize streamlines as valid or invalid connections.

Candidate bundles are extracted in the same way that we defined the gold standard bundles. First, we run RBX to extract candidate bundles from the candidate whole-brain tractogram. Candidate bundles are then converted to binary volume coverage masks. Finally, each candidate mask is compared against its corresponding gold standard bundle mask to compute evaluation metrics.

For each subject in the testset, and for each available bundle of the given subject, we extract the following evaluation metrics: Dice score, overlap and overreach. The scores are averaged over all subjects of the testset to provide final scores. Altogether, these metrics help better understand the performance of a candidate tractography algorithm.

The evaluation pipeline is available online (https://github.com/scil-vital/TractoInferno/) and should be used with the provided *TractoInferno* test-set, along with the default RBX-flow models.

## 5. RNN-based tractography

To gauge the performances of ML models trained on the *TractoInferno* dataset, we implemented an RNN model and the necessary framework to train it on a large-scale tractography database, which was used multiple times in published papers in the last few years, such as *Learn2Track* [18], *DeepTract* [22], and *Entrack* [20]. Using the base implementation, we can easily modify the last layer of the model and its loss function to mimic the mentioned RNN models, and a few more.

We choose the stacked Long Short-Term Memory (LSTM) network as the recurrent building block for conditional streamline prediction. The LSTM is a type of RNN designed specifically to handle long-term dependencies, with the ability to deal with exploding and vanishing gradient problems [24].

### 5.1. Learn2track

*Learn2track* [18] proposed an RNN model for tractography, where the out-put of the model at each timestep is a 3D vector, used as the next direction of the streamline. The predicted vector is then scaled to the chosen step size, in order to match the lengths of the target and prediction.

From the same idea, we implemented an LSTM for deterministic tractography. As in the original *learn2track* paper, we used the squared error loss function between the target and prediction. The loss for a single streamline *S* composed of *T* steps is the following squared error:

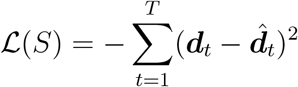

where ***d***_*t*_ and 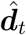 are the target and predicted directions. This model is noted as *Det-SE*.

However, to accurately reflect that only the direction of the predicted vector is important (not the magnitude), we also performed an experiment where we minimized the negative cosine similarity between the target and predicted directions:

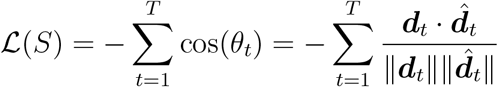

where *θ_t_* is the angle between ***d***_*t*_ and 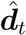. This model is noted as *Det-Cosine*.

### 5.2. DeepTract

In the same spirit as *learn2track*, *DeepTract* [22] is a recurrent model for probabilistic tractography. In this case, the model output is a distribution over classes, where each class corresponds to a direction on the unit sphere, i.e. a discrete conditional fiber ODF.

As in the original paper, we implemented a cross-entropy loss function:

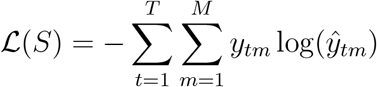

where *M* is the number of classes, and *y_t_* and 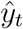 are vectors of target and predicted class probabilities. Note that we did not use label smoothing as in the original paper, nor entropy-based tracking termination. This model is noted as *Prob-Sphere*.

### 5.3. Entrack

*Entrack* [20] is a non-recurrent artificial neural network for probabilistic tractography. The model is instead a feed-forward neural network, but includes the previous streamline direction as prior information to guide the tracking process. The model outputs the parameters for a von Mises-Fisher distribution, i.e. a 3D unit-length vector for the mean, and a scalar concentration parameter. The distribution is analogous to a Gaussian distribution, but defined on the unit sphere instead of euclidean space.

We chose to apply the same general idea, using a recurrent network that predicts the parameters for a von Mises-Fisher distribution on a 3D sphere.

We used the negative log-likelihood of the von Mises-Fisher distribution as the loss function:

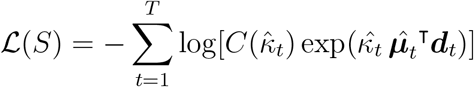

where the predicted parameters of the distribution are 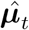 (a unit-length vector) and 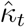 (a scalar concentration parameter), and ***d***_*t*_ is the target unit-length vector at step *t*. 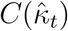 abbreviates the normalization constant associated with the distribution, defined as following in the 3-dimensional case:

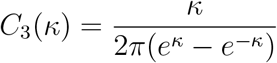

Note that unlike the original method, we didn’t use an entropy maximization scheme to regularize the predicted distribution. This implementation is noted as *Prob-vMF*.

### 5.4. Gaussian distribution output

Following *Entrack* and the idea of predicting the parameters of a continuous probability distribution, we implemented another model, using a multivariate Gaussian distribution instead of a von Mises-Fisher distribution. This model outputs a 3D vector for the mean, and 3 scalars for the variance, (one in each dimension). We choose to use a diagonal covariance matrix, for stability, and do not output any values for covariance.

In the 3-dimensional case, the negative log-likelihood loss function is:

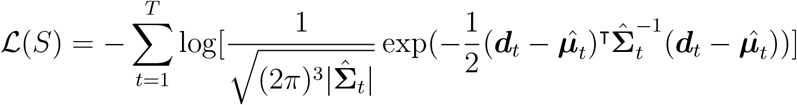

where 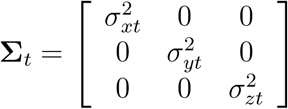 is the predicted diagonal covariance matrix at streamline step *t*. This model is noted as *Prob-Gaussian*.

### 5.5. Gaussian mixture distribution output

The previous Gaussian model outputs a single average direction which is appropriated in most cases. However, there may be cases of bundle fanning or forking where the single-mode assumption may be an issue. This is because the Gaussian probability density can only be spread over a large area.

As such, some regions may be better modelled with more than one location of higher density. To this end, we implemented a mixture density network[64] using a mixture of 3 Gaussian distributions. For each Gaussian, the model outputs 1 mixture weight, a 3D vector for the mean, and 3 scalars for the variances (again, we fix the covariances to zero).

In the 3-dimensional case, using a mixture of 3 Gaussians, the negative log-likelihood loss function is:

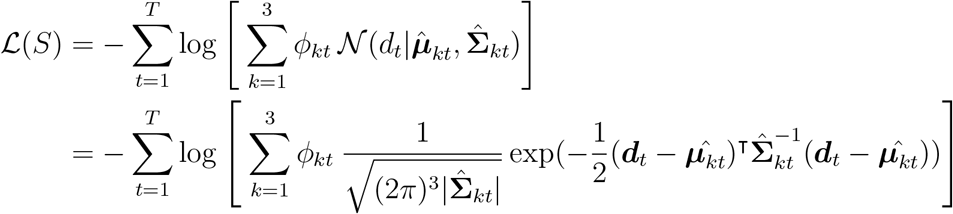

where *k* denotes the number of Gaussians in the mixture, and *ϕ_kt_* is the mixture parameter for the Gaussian *k* at streamline step *t*. This model is noted as *Prob-Mixture*.

### 5.6. Implementation details

All models were composed of 5 hidden layers of 500 units, used dropout with a rate of 0.1, and a batch size of 50 000 streamline steps. We added skip connections from the input layer to all hidden layers, and from all hidden layers to the output layer, inspired by [65]. We applied layer normalization [66] between all hidden layers, in order to stabilize the hidden state dynamics in recurrent neural networks. We used the Adam optimizer with the default parameters.

For all experiments, we used the maximal spherical harmonics (SH) coefficients of order 6 fitted to the TractoFlow-processed DWI signal as the input signal. In all cases, the models were trained using the exact same training/validation/test datasets, with a streamline step size fixed to 1.0 mm for training and tracking. To help guide the model, we also included as input the diffusion signal in a neighbourhood of 6 directions (two for each axis, positive and negative) at a distance of 1.2 mm.

All models were trained for a maximum of 30 epochs (corresponding to around 2 weeks of training time on a 16Gb NVidia V100SXM2), but early stopping was used to stop training when the loss has not improved after 5 epochs. Each epoch was capped to 10 000 updates, as the sheer size of the dataset would otherwise require multiple days of training for a single epoch.

## 6. Results & Discussion

We report in Table 4 the results of the *TractoInferno* evaluation pipeline for each individual tractography algorithm used to build the reference bundles, and for every model detailed in Section 5 after the training procedure.

**Table 4:** Tractography evaluation results on the *TractoInferno* dataset. The Prob-vMF model did not produce any results, and is noted as {N/A}.

|  | Dice | Overlap | Overreach |
| --- | --- | --- | --- |
| *Reference methods* |  |  |  |
| Deterministic | 0.397 | 0.267 | 0.029 |
| Probabilistic | 0.553 | 0.433 | 0.068 |
| PFT | 0.680 | 0.688 | 0.266 |
| SET | 0.624 | 0.570 | 0.184 |
| Ensemble (Det+Prob+PFT+SET) | 1.000 | 1.000 | 0.000 |
| *RNN-based methods* |  |  |  |
| Det-SE (Learn2track) | 0.580 | 0.495 | 0.172 |
| Det-Cosine | 0.606 | 0.535 | 0.204 |
| Prob-Sphere (DeepTract) | 0.601 | 0.534 | 0.202 |
| Prob-vMF (Entrack) | N/A | N/A | N/A |
| Prob-Gaussian | 0.624 | 0.585 | 0.264 |
| Prob-Mixture | 0.407 | 0.284 | 0.053 |

Of all the base algorithms used to build the reference tractograms, PFT performed the best in terms of Dice score and overlap. This is consistent with the fact that it is a state-of-the-art algorithm, and works best when trying to fill the space with streamlines. However, we show that no algorithm can single-handedly account for the gold standard, and using the union of all methods provides a more complete reconstruction.

In both traditional and RNN-based variants, models with the best Dice/ overlap results also had the worst overreach score. However, in the case of bundle reconstruction, it is less of a concern, because there is always a possibility of applying post-processing techniques to filter streamlines. Also, since our gold standard is not perfect, it might not cover the whole possible space as delineated by the RBX algorithm. Furthermore, because the scores are evaluated using binary bundle masks, a small number of streamlines can easily cross a high number of overreaching voxels. Ultimately, the goal is to find a model that can cover as much space as possible, so the overreach score is an interesting information to have, but is not the best indicator of performance in our case.

Of all the RNN-based methods, the Gaussian output model obtained the best Dice score and overlap, hinting that a probabilistic model works best. This is in line with traditional probabilistic algorithms being more suited to bundle reconstruction than deterministic approaches.

Given the worse performance of other probabilistic models, it seems that adding complexity is not beneficial. Training an RNN with a more complex distribution like the mixture of Gaussians might require a different architecture, or more model capacity, to achieve better results. Unfortunately, the RNN with a von Mises-Fisher output had a hard time training, and produced erratic streamlines that mostly did not survive the evaluation pipeline. It would seem that training the vMF distribution is too unstable when using a likelihood loss function, and performing an entropy maximization procedure like the original authors might be required to have a stable training procedure.

Across all results (both reference algorithms and RNN-based methods), the general trend holds that with a better Dice score and overlap, there is also more overreach. This indicates that there is still work to be done to limit the production of false positive streamlines.

To illustrate the differences between algorithms, we showcase the reconstructions of three bundles taken from a random test subject. We chose bundles of both *medium* and *hard* difficulty for tractography, as reported in [2]. Figure 3 shows a part of the *Corpus Callosum* (medium difficulty), while Figures 4 and 5 show the *Optic Radiation* and the *Pyramidal Tract* (hard difficulty). Note that in all cases, as mentioned before, the Prob-vMF method did not produce any meaningful results, which explains why no results are shown.

**Figure 3:**
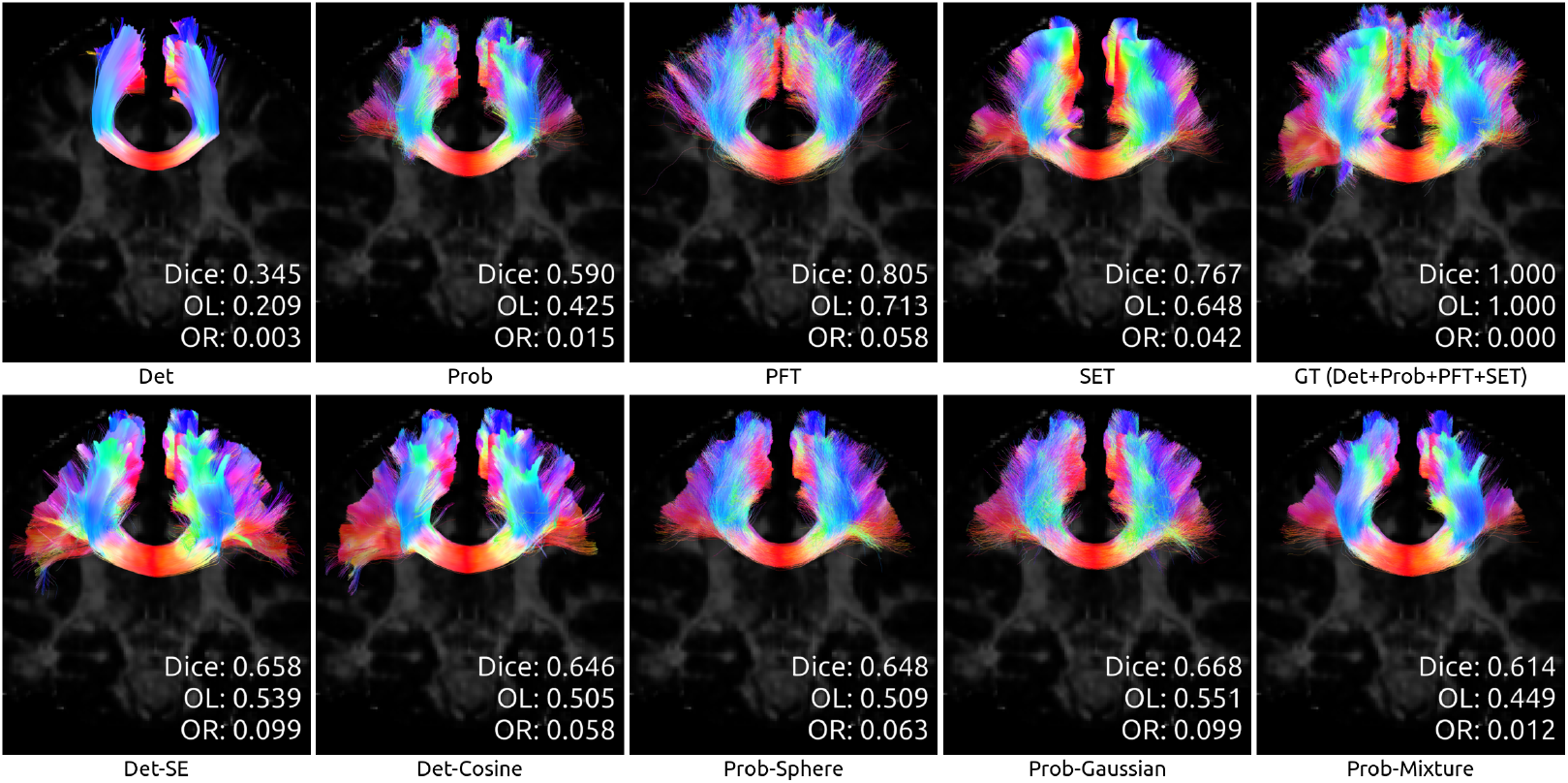
Reconstruction of the *Corpus Callosum* (medium difficulty) by all algorithms, for test subject sub-1006.

**Figure 4:**
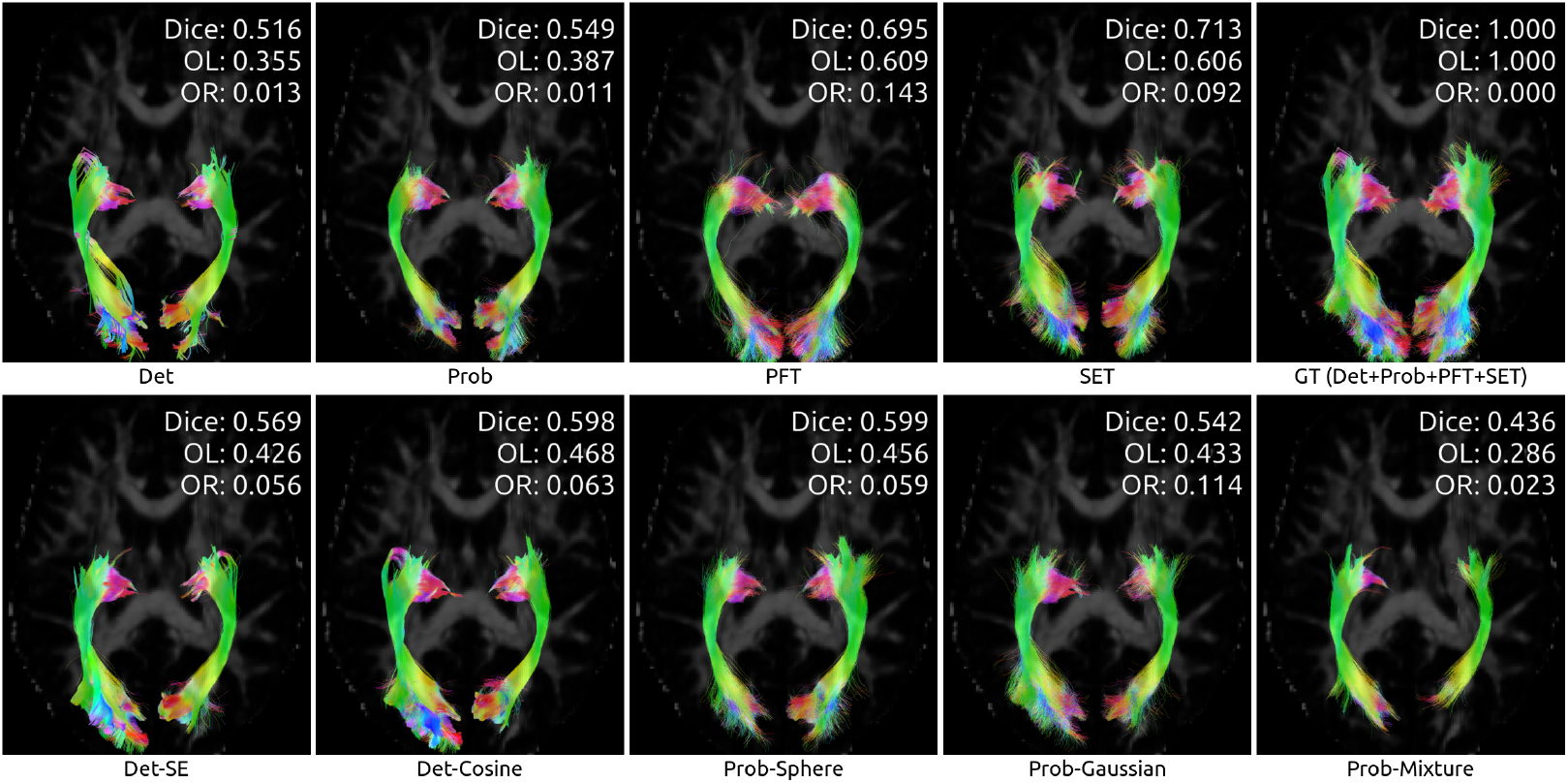
Reconstruction of the *Optic Radiation* (hard difficulty) by all algorithms, for test subject sub-1006.

**Figure 5:**
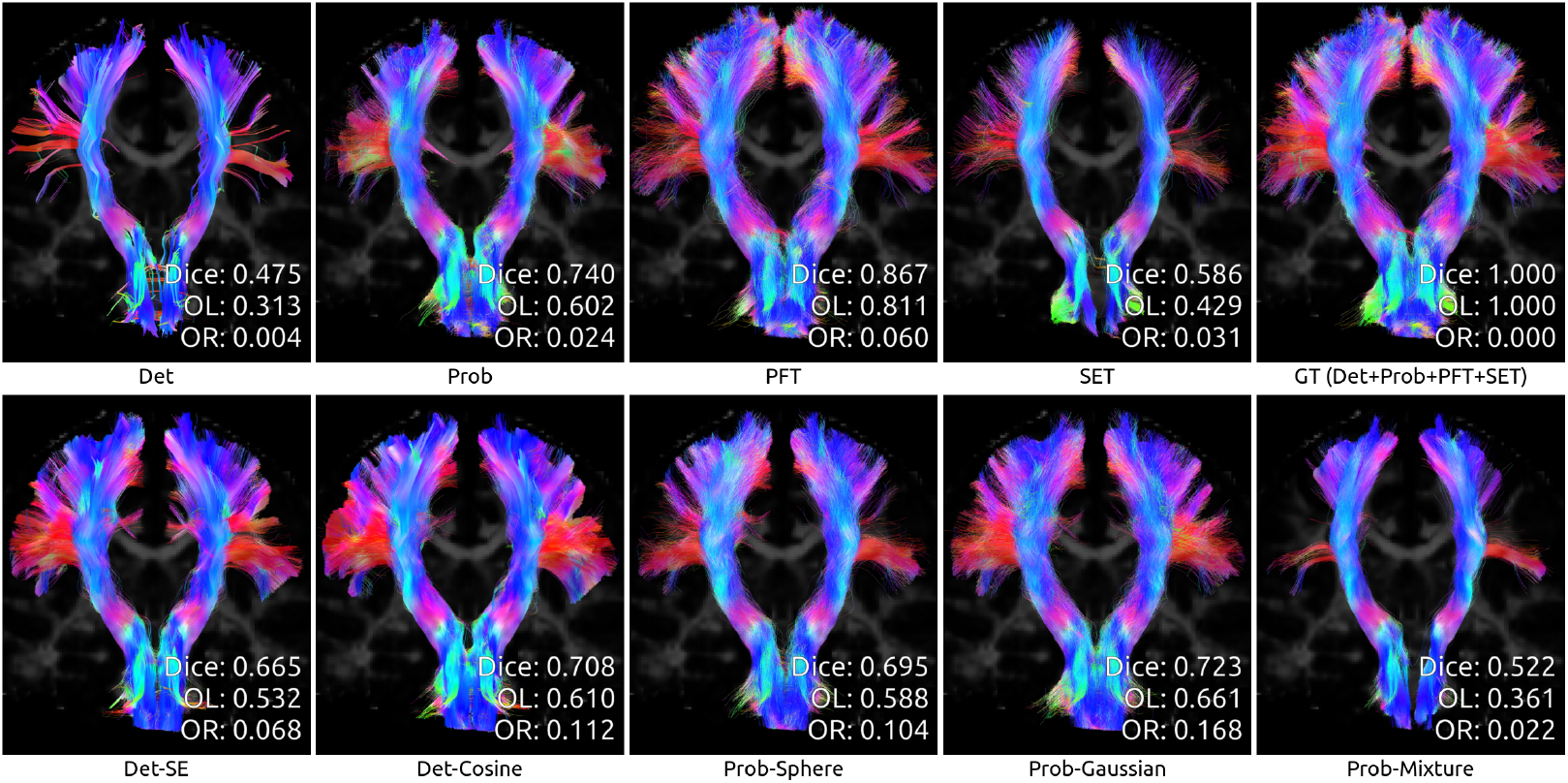
Reconstruction of the *Pyramidal Tract* (hard difficulty) by all algorithms, for test subject sub-1006.

Also of note, RNN-based models seem to get results on par with traditional algorithms, but not quite as good as the state-of-the-art Particle-Filtered Tractography. However, Poulin et al. produced results far beyond even PFT using an RNN approach trained on a single-database, using a single-bundle per model [67]. While we did not train any model with the single-bundle approach on *TractoInferno*, both results hint that there is a need for more data, more model capacity, or for specialization of algorithms, in order to outperform currently-used methods. We advocate that *TractoInferno* is one way to investigate this problem further.

## 7. Conclusion

We provide an open-access, multi-site dMRI and tractography database aimed at training and evaluating machine learning tractography models. It combines data from multiple datasets, and applies the same processing and QC steps for a uniform database. We also produce results using the available evaluation pipeline for both traditional algorithms and machine learning models based on a recurrent architecture.

We offer *TractoInferno* as a solution to the multiple issues already reported in the literature for machine learning tractography. Indeed, while such algorithms have been proposed in the last few years with promising results, none has been shown to be the fundamental solution to classical tractography. They commonly suffer from variable training data, dissimilar evaluation method, and limited dataset size, among others. To this end, a uniform, large-scale, and multi-site database such as *TractoInferno* is an essential tool, paving the way for reproducible and comparable research among machine learning tractography researchers.

## 8. Data access

The *TractoInferno* database is freely available online on the OpenNeuro platform: https://openneuro.org/datasets/ds003900. The evaluation pipeline is avalaible on GitHub: https://github.com/scil-vital/TractoInferno/.

## 9. Special thanks

Thanks to Laurent Petit for his help with the BIL & GIN database.

## 10. Acknowledgements

P.P. was supported by the FQRNT, grant #206270. This research was enabled in part by by Compute Canada (www.computecanada.ca) and CBRAIN [62].

## Appendix A. *TractoInferno* pipeline execution commands

All commands used to process the *TractoInferno* dataset are reported here. The input files and directories for each command might need to be reorganized between steps; refer to the specific package documentation for more details.

## Appendix A.1. QC DWI

URLs:

- https://github.com/scilus/dmriqcpy
- https://github.com/scilus/dmriqc_flow

Command:

~~~
nextflow run dmriqc-flow-0.1.2/main.nf -profile input_qc
--root input/
-with-singularity singularity_dmriqc_0.1.2.img -resume
--raw_dwi_nb_threads 10
~~~

## Appendix A.2. TractoFlow

URL: https://github.com/scilus/tractoflow/

Command:

~~~
nextflow run tractoflow-2.1.1/main.nf --root input/
--dti_shells "0 700 1000 1200"
--fodf_shells "0 700 1000 1200"
-with-singularity tractoflow_2.1.1_650f776_2020-07-15.img
-resume -profile fully_reproducible --mean_frf false
--set_frf true --nbr_seeds 1
~~~

## Appendix A.3. QC TractoFlow

URLs:

- https://github.com/scilus/dmriqcpy
- https://github.com/scilus/dmriqc_flow

Command:

~~~
nextflow run dmriqc-flow-0.1.2/main.nf
-profile tractoflow_qc_light
--root/ ../TractoFlow/results
-with-singularity singularity_dmriqc_0.1.2.img -resume
~~~

## Appendix A.4. SH signal fitting

URL: https://github.com/ppoulin91/tractoinferno_compute_sh_flow

Command:

~~~
nextflow run code/tractoinferno_compute_sh_flow/main.nf
--input input/ --use_attenuation --sh_order 6
-with-singularity scilus-1.2.1.img -resume
~~~

## Appendix A.5. Tractography (Det, Prob, PFT)

URL: https://github.com/ppoulin91/tractoinferno_tracking_flow

Command:

~~~
nextflow run code/tractoinferno_tracking_flow/main.nf
--input ../TractoFlow/results
-iwith-singularity tractoflow_2.1.1_650f776_2020-07-15.img
-iresume
~~~

## Appendix A.6. Tractography (SET)

URLs:

- https://github.com/StongeEtienne/set-nf
- https://github.com/scilus/convert_set_flow

Commands:

~~~
nextflow run code/set-nf/main.nf
--tractoflow ../TractoFlow/results
--surfaces ../civet/results -profile civet2_dkt
-with-singularity set_1v1.img -resume
nextflow run code/convert_set_flow/main.nf
--root_set ../SET/results
--root_tractoflow ../TractoFlow/results
-with-singularity scilus-1.2.1.img
-resume
~~~

## Appendix A.7. RecoBundlesX (RBX)

URL: https://github.com/scilus/rbx_flow/

Command:

~~~
nextflow run code/rbx_flow/main.nf -resume
-with-singularity scilus-1.2.0_rbxflow-1.1.0.img
-profile large_dataset --input input/
--atlas_config code/rbx-atlas/config.json
--atlas_anat code/rbx-atlas/mni_masked.nii.gz
--atlas_directory code/rbx-atlas/atlas/
--run_average_bundles false
~~~

## Appendix A.8. QC RBX

URLs:

- https://github.com/scilus/dmriqcpy
- https://github.com/scilus/dmriqc_flow

Command:

~~~
nextflow run code/dmriqc_flow/main.nf -resume
-with-singularity
singularity_dmriqcflow_hotfix_scilpy_1.2.0.img
-profile rbx_qc --input ../RBX/results/
~~~

